# Disruption of Redox Balance Enhances the Effects of BRAF-inhibition in Melanoma Cells

**DOI:** 10.1101/818989

**Authors:** B. Bishal Paudel, Joshua E. Lewis, Keisha N. Hardeman, Corey E. Hayford, Charles J. Robbins, Simona G. Codreanu, Stacy D. Sherrod, John A. McLean, Melissa L. Kemp, Vito Quaranta

**Affiliations:** Department of Biochemistry, Vanderbilt University, Nashville, TN.; Quantitative Systems Biology Center (QSBC), Vanderbilt University, Nashville, TN.; Department of Biomedical Engineering, University of Virginia, Charlottesville, VA.; The Wallace H. Coulter Department of Biomedical Engineering, Georgia Institute of Technology and Emory University, Atlanta, Georgia; Chemical and Physical Biology Graduate Program, Vanderbilt University, Nashville, TN.; Center for Innovative Technology, Department of Chemistry, Vanderbilt Institute of Chemical Biology, Vanderbilt University, Nashville, TN.

**Keywords:** Systems biology, Redox capacity, Drug insensitivity, Combination targets, Antioxidants

## Abstract

**Summary:** Melanomas harboring *BRAF* mutations can be treated with BRAF inhibitors (BRAFi), but responses are varied and tumor recurrence is inevitable. Here, using an integrative approach of experimentation and mathematical flux balance analyses in *BRAF*-mutated melanoma cells, we report that elevated antioxidant capacity is linked to BRAFi sensitivity in melanoma cells. High levels of antioxidant metabolites in cells with reduced BRAFi sensitivity confirm this conclusion. By extending our analyses to other melanoma subtypes in TCGA, we predict that elevated redox capacity is a general feature of melanomas, not previously observed. We propose that redox vulnerabilities could be exploited for therapeutic benefits and identify unsuspected combination targets to enhance the effects of BRAFi in any melanoma, regardless of mutational status.

## Introduction

Targeted therapy has been a major breakthrough for melanoma patients harboring *BRAF*^V600^ mutations because of a superior response rate, and a remarkable short-term efficacy (Chapman et al., 2011; Sosman et al., 2012). However, the clinical responses are highly variable and short-lived, and relapse is almost universal (Shi et al., 2014; Sosman et al., 2012). Overcoming reduced sensitivity and acquired resistance to targeted therapy is a major goal of current melanoma research. Several mechanisms of reduced sensitivity have been proposed (Poulikakos et al., 2011; Shi et al., 2014; Thompson et al., 2014; Wagle et al., 2011), which have led to the development of several combination regimens in melanoma either with other targeted therapies (Flaherty et al., 2012; Larkin et al., 2014; Menzies and Long, 2014), or in conjunction with immunotherapies (Hu-Lieskovan et al., 2015). While these therapies improve responses, treatment outcomes still vary, and benefits remain transient and unpredictable (Luke et al., 2017).

Some resistance can be attributed to genetic mutations (Greaves and Maley, 2012; Nowell, 1976), but accumulating evidence indicates that nongenetic processes play a critical role in response of cancer cells to drug treatment (Niepel et al., 2009; Paudel et al., 2018; Shaffer et al., 2017; Smith et al., 2016). Metabolic reprogramming, recognized as a hallmark of cancer, has recently emerged as a potential nongenetic process that contributes to the emergence of drug-tolerant cancer cells (Hanahan and Weinberg, 2011). Otto Warburg first reported a link between tumor and metabolism in his influential observation that cancer cells convert most intracellular glucose to lactate--that is aerobic glycolysis (Warburg, 1925). Recent studies have shown that cancer cells can utilize glycolysis, mitochondrial respiration, or both, depending on their environment (Dang, 2012). Furthermore, cancer cells can adapt metabolically in response to external perturbations (DeBerardinis et al., 2008; Jia et al., 2019). This metabolic flexibility provides cancer cells with energy, and necessary intermediates for biosynthetic processes required for survival and to maintain redox balance under changing environments (DeBerardinis et al., 2008; Paudel and Quaranta, 2019). Furthermore, metabolic pathways are complex and interconnected, warranting a systems level approach to examine their relative importance, and how they change in cancer cells. Flux Balance Analysis (FBA) is the most commonly used mathematical modeling approach for genome-scale metabolic network reconstructions to estimate the role of metabolic reactions in a network (Blazier and Papin, 2012; Orth et al., 2010). Such a quantitative approach can be utilized to predict global metabolic states of cancer cells under various conditions.

Melanoma cells upon BRAF-inhibition have been shown to induce an enhanced oxidative phosphorylation (Delgado-Goni et al., 2016; Haq et al., 2013; Smith et al., 2016). Byproducts of augmented mitochondrial activity are reactive oxygen species (ROS), and this metabolic switch promotes oxidative stress in cells (Cesi et al., 2017; Zorov et al., 2014). However, the cellular response to ROS is rather complex: low levels facilitate intracellular signaling, while high levels may cause cell death (Dikalov et al., 2011; Panieri and Santoro, 2016; Schieber and Chandel, 2014). Therefore, cells require a robust antioxidant defense system to respond to an accumulation of ROS. By invoking antioxidant systems, cancer cells can clear excess ROS levels within a range that is not detrimental to them--a high redox homeostatic state (DeNicola et al., 2011). Nicotinamide adenine dinucleotide phosphate (NADPH), and glutathione (GSH) are two major antioxidants that maintain redox homeostasis in cells (Panieri and Santoro, 2016). NADPH is produced via the Pentose Phosphate Pathway (PPP), and acts as a shared substrate for both GSH regeneration, and ROS production, thus maintaining an optimum redox balance within cancer cells. Even within cancer subtypes, this redox state could be heterogeneous (Sarmiento-Salinas et al., 2019), motivating further research to examine how this balance could be altered for therapeutic benefits.

The link between oxidative stress and drug response in melanoma is beginning to be explored (Mishra et al., 2018; Zaal and Berkers, 2018). BRAF-inhibitor resistant melanomas were shown to upregulate *NRF2*-mediated antioxidant response to maintain cell survival (Khamari et al., 2018). Nonetheless, it still remains to be examined how redox potential of melanoma cells affects their drug sensitivity, and how it is maintained under BRAF-inhibition. These are important considerations, raising the possibility that redox balance can be modulated to enhance the effects of existing therapies (Yuan et al., 2018).

Here, we show that antioxidant capacity of melanoma cells is linked to their drug sensitivity. Using an integrative approach through bioinformatics and FBA, we show that melanoma cells with reduced sensitivity to BRAFi exhibit an enhanced capacity for anti-oxidation and redox buffer, specifically through NADPH oxidizing enzymes. By directly quantifying the redox-related metabolites, we confirm that drug-insensitive melanoma cells can maintain higher levels of antioxidant metabolites during treatment. Furthermore, we report that pharmacological disruption of redox axis involving glutathione enhances the effects of targeted therapies. We also extended our analysis to other melanoma subtypes in The Cancer Genome Atlas (TCGA), and found that elevated redox capacity could be a general feature of melanoma. Our results thus provide a proof-of-principle that redox vulnerabilities could be exploited for therapeutic benefits in melanoma.

## Results

### Gene expressions and Flux Balance Analyses reveal enhanced capacity of redox balance in cells with reduced sensitivity to BRAF-inhibition

We recently reported that *BRAF*-mutated melanoma cell lines, including isogenic single-cell derived subclones, exhibit varying drug sensitivities to a small molecule *BRAF* kinase inhibitor (BRAFi) (Hardeman et al., 2017; Paudel et al., 2018). Using Drug-Induced Proliferation (DIP) rates (***Table S1***) (Harris et al., 2016), as a measure of drug effects, we examined the molecular correlates of *BRAF*i sensitivity, and reported that top 200 differentially expressed genes (DEGs) in drug-insensitive cells were enriched in processes and functions related to redox metabolism (Meyer et al., 2019). Here, we extended our analysis to examine all significant DEGs among isogenic subclones (***Table S2***) using their RNASeq profiles. Among 2,165 DEGs, 1361 (⅔) were up-regulated, while 804 (⅓) were down-regulated (***Fig. 1A, B***). Next, we projected all DEGs onto a larger panel of *BRAF*-mutated melanoma cells from CCLE datasets (Barretina et al., 2012), and calculated their correlation to DIP rates. We selected significantly correlated genes using a criterion as outlined in ***Fig. 1A***, and identified both positively and negatively correlated genes (***Fig. 1C*** ***& Table S3***). Gene Ontology (GO) analysis on positively correlated genes in both subclones (**Fig. 1B**), and CCLE melanoma panel (**Fig. 1C**) showed enrichment of molecular functions related to redox balance, coenzyme metabolic process, and oxidoreductase activity (***Fig. 1D*** ***& Table S4***). Specifically, we observed enrichment of gene signatures (CYBA, NADPH Oxidase 5 (NOX5), HTATIP2, SLC7A11, highlighted in magenta in **Fig. 1B** **&** **C**) related to redox balance and modulation, oxidoreductase activity, and reactions that utilize or consume antioxidant NADPH as a substrate (***Fig. 1E***). To quantify the contribution of redox enzymes toward NADPH oxidation, we next predicted steady-state fluxes through several major NADPH-oxidizing reactions using a previously published framework for integrating transcriptomic, kinetic, and thermodynamic data into Flux Balance Analysis (FBA) models of individual cancer cell lines (Lewis et al., 2018; Orth et al., 2010). The drug-insensitive SC10 model showed significantly greater total conversion of NADPH to NADP+ compared to the drug-sensitive SC01 model. While most reactions did not display significant differences in predicted flux distributions, the SC10 model had significantly higher fluxes through Glutathione Disulfide Reductase (GSR), dihydrofolate reductase (DHFR), and NADPH oxidase (NOX1/3/5/CYBB) (***Fig. 1F*** ***& Table S5A***). Consistent with the results in isogenic subclones, the subset of CCLE melanoma cell lines which are insensitive to MAPK-pathway inhibition also had higher predicted fluxes through GSR and DHFR reactions (***Fig. 1G*** ***& Table S5B***). Taken together, these results suggest that melanoma cells with reduced drug sensitivity, as measured by DIP rates, exhibit enhanced anti-oxidation and redox balance. Specifically, we found that NADPH oxidation reactions coupled to glutathione exhibited higher gene expression and predicted fluxes in drug-insensitive melanoma cells.

**Figure 1:**
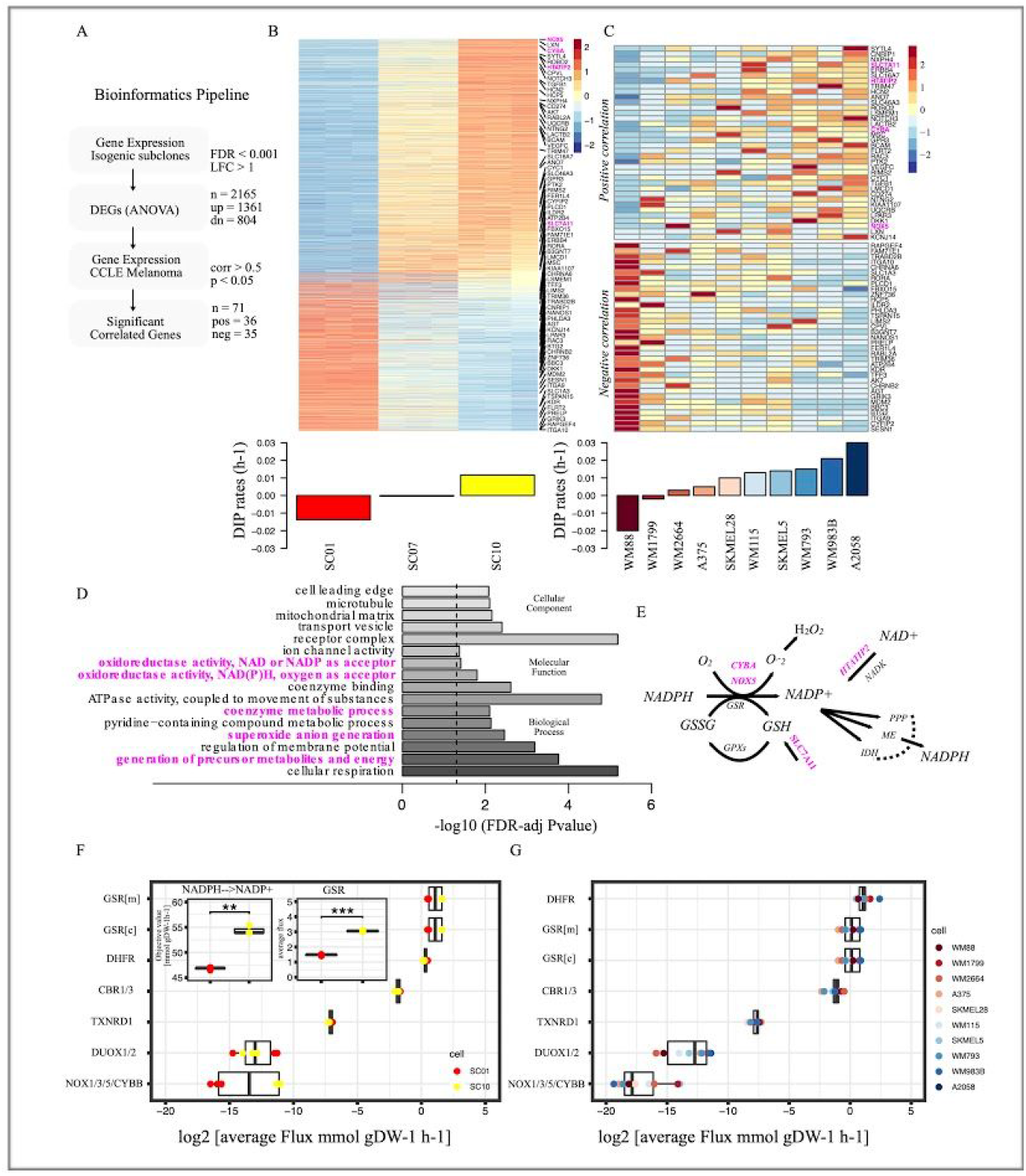
*BRAF*-mutated melanoma cells with reduced sensitivity to BRAF-inhibition show elevated redox capacity. (A) Bioinformatics pipeline to identify genes correlated with drug sensitivity, extension of an approach in previously published report (Meyer Pt al.. 2019) Briefly, Differentially Expressed Genes (DEGs) were identified from gene expression of isogenic subclones. Expressions of the identified genes were then correlated with DIP rates in CCLE melanoma panel. Statistical criteria used shown in the pipeline. (B) Heatmap of DEGs among subclones (top); Barplot of DIP rates of three subclones, SC01, SC07, and SC10 in 8μM PLX4720 (*bottom*); (C) Heatmap of 71 genes from melanoma cell lines in COLE panel that show a significant correlation between their expression and drug sensitivity. Genes in upper panel show positive correlation, while genes in lower panel show negative correlation to DIP rates. Highlighted in magenta are the redox-related genes involved in NADPH oxidation (*top*); Barplot of quantified DIP rates for the indicated melanoma cells in 8μM PLX4720 (*bottom*). (ID) Enriched Gene Ontology (GO) terms of the significant, and positively correlated genes from Figs. 1 B, and C, and their FDR-adjusted p-values on barplot. (E) Schematic of the redox axis of NADPH oxidation, genes highlighted in magenta are the genes that are positively correlated in Fig. 1C. (F) Average FBA predicted metabolic fluxes through NADPH oxidation reactions between two single-cell-derived SKMEL5 subclones models, SC01, and SC10; three replicates of the model were considered. Inset (left) shows the objective value for net conversion of NADPH to NADP+ for two subclones; inset (right) average fluxes through one of NADPH oxidizing reactions, GSR. (H) Average FBA predicted metabolic fluxes through NADPH oxidation reactions in CCLE melanoma cell line models.

### Redox metabolites with antioxidant properties are maintained at higher levels after BRAF-inhibition in melanoma cells with reduced sensitivity to BRAF-inhibition

To directly assess the metabolite abundance, we performed a combined global untargeted-targeted liquid chromatography-mass spectrometry (LC-MS/MS) metabolomics analysis in *BRAF*-mutated melanoma cells under *BRAF*-inhibition. In particular, we selected one drug-sensitive (WM88), and one drug-insensitive (SC10) melanoma cell line, and compared their metabolite profiles both in the absence and presence of *BRAF*-inhibitors (***Table S6***). In total, 2687 features or metabolites (unique retention time and m/z pairs) were detected, quantified and analyzed in these studies. Unsupervised Principal Component Analysis across all detected metabolites clustered sensitive melanoma cells away from insensitive cells at baseline along principal component 1 (PC1), while the drug-treated samples are separated along principal component 2 (PC2) (***Fig. 2A***). In general, the cells exhibit distinct metabolite profiles, with BRAF-inhibition inducing a marked metabolite profile shift (***Fig. 2A, B***). In these studies, 18 validated (Level 1) redox metabolites were quantified. These metabolites were validated through accurate mass measurements (<5 ppm error), similarities in isotope distribution (>90%), matched retention times, and matched fragmentation spectra based on in-house databases (Schrimpe-Rutledge et al., 2016). In sum, we observed 2687 metabolites and using the combined global metabolomic profiles were able to mine the data for high confident metabolite identification for molecules of interest (redox, GSH synthesis and energy metabolites).

**Figure 2:**
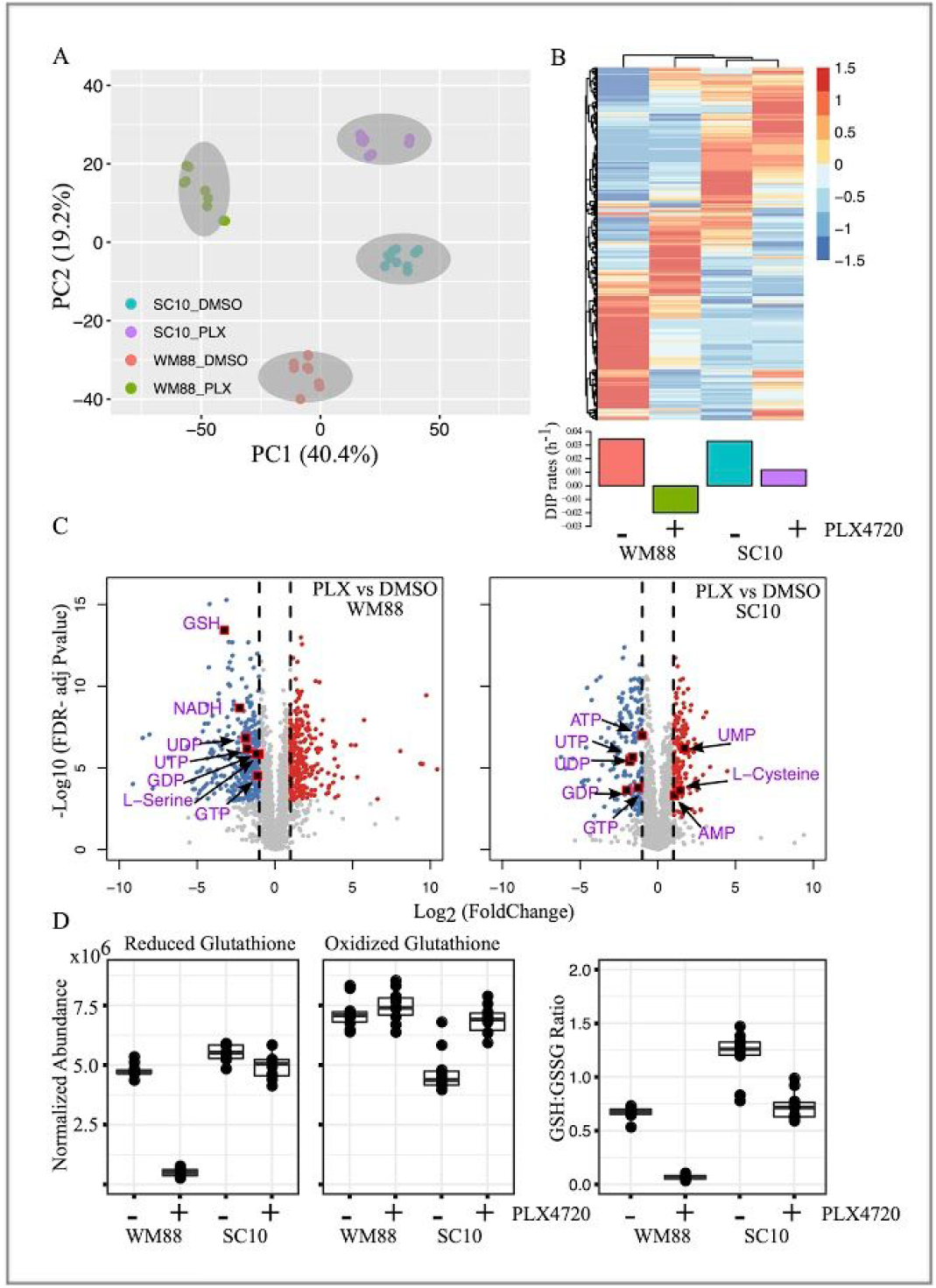
Global metabolomic analysis reveals distinct metabolite profiles in BRAF-mutated melanoma cells with differing drug sensitivities. (A) Principal component analysis (PCA) of metabolite profiles for two BRAF-mutated melanoma cells, WM88, and SC10 (SKMEL5-SC10), treated either in DMSO or 8μM PLX4720 for 24 hrs. The two cell lines cluster and separate along PC1, while the drug-treated samples separate along PC2. (b) Heatmap analysis of identified metabolites in two melanoma cells (WM88 and SC10) either in DMSO or PLX4720 treatment (top) provides a global comparison of the relative abundances of individual metabolite compounds across the different groups; Barplot of DIP rates in WM88, and SC10 in DMSO or in 8μM PLX4720 (*bottom*). (C) Volcano plots showing differentially expressed metabolites after treatment of two melanoma cells (WM88 or SC10) with 8μM PLX4720 for 24 h. The negative log10 transformed Bonferroni corrected P-values (y-axis) are plotted against the average log2 fold changes in metabolite abundance (x-axis). Grey represents metabolites not differentially expressed, red are upregulated metabolites, and blue are downregulated metabolites. Vertical dotted lines represent log2 fold change of 1 or −1. Statistical cutoff of FDR-adjusted p-values < 0.001, and log-fold change greater or equal to 1 was used. (D) Box plots showing normalized abundances of reduced glutathione (GSH), oxidized glutathione (GSSG), and the ratio of GSH and GSSG in indicated melanoma cells treated with either DMSO or 8μM PLX4720 for 24 h: the solid line is the median, the box spans the first, and third quartiles, the whiskers extend to 1.5× the interquartile range--total of 10 replicates shown.

To probe the treatment-dependent changes in metabolite abundances, we also performed pairwise comparisons between BRAFi treated and control samples within each cell line. In particular, we focused on the redox-related metabolites based on our earlier results on anti-oxidation and redox balance. Differentially expressed metabolites were determined based on a statistical cutoff of FDR-adjusted p-values < 0.001, and log-fold change greater or equal to 1. PLX4720 treatment significantly depleted the levels of energy metabolites such as GTP, UTP, ATP, GDP in both cell lines (***Fig. 2C***, ***Fig. S1A***). Among the redox metabolites, there was an increase in monophosphate nucleotides such as UMP, AMP in SC10 (***Fig. 2C***). Interestingly, reduced GSH levels were significantly depleted only in WM88, and not in SC10 upon BRAFi treatment (***Fig. 2C***). GSH levels, however, were comparable between two cells at baseline (***Fig. 2D***, ***Fig. S1B***). Because glutathione can exist in both reduced (GSH), and oxidized form (GSSG), we also quantified the levels of GSSG in both WM88 and SC10 cells. GSSG levels were lower in SC10 compared to WM88 at baseline, but similar in PLX4720 treatment (***Fig. 2D***). The ratio of reduced and oxidized glutathione, often used as an indicator of oxidative stress in cells, were also calculated. These data show that SC10 cells had a significantly higher GSH:GSSG ratio at baseline compared to drug-sensitive WM88 cells, rendering them a robust antioxidant capacity (***Fig. 2D***). Upon treatment, SC10 showed a slight decrease in GSH:GSSG ratio (μ_DMSO = 1.20, μ_PLX = 0.732, decrease of 1.6 fold). In contrast, WM88 showed an approximately 10-fold decrease in its ratio upon BRAF-inhibition (μ_DMSO = 0.67, μ_PLX = 0.067) (***Fig. S1C***).

Additionally, we also quantified the abundance of cysteine, cystine, and Serine--all precursors of *de novo* GSH synthesis. Transport of cystine into cells to form cysteine depends on the activity of cystine/glutamate antiporter, SLC7A11, one of the genes positively correlated with DIP rates. At baseline, we did not detect cystine, therefore, we quantified the ratio of cystine and cysteine in drug-treated conditions. The ratio of cystine and cysteine under treatment was significantly higher in WM88 than in SC10, indicating a rapid accumulation of extracellular cystine (***Fig. S1D***). Additionally, BRAFi treatment significantly increased the levels of Cysteine in SC10, and not in WM88 (***Fig. 2C***). Levels of serine were substantially higher in SC10 compared to WM88 at baseline, and upon BRAFi treatment were maintained in SC10, and significantly decreased in WM88 (***Fig. 2C***, ***Fig. S1A, S1E***). Collectively, these results indicate that drug-insensitive cells have a robust anti-oxidation and redox system that can be maintained under BRAF-inhibition, perhaps through both an efficient regeneration and *de novo* synthesis of GSH.

### Pharmacological inhibition of redox enzymes synergizes with MAPK-inhibition in BRAF-mutated melanoma cells

Based on our results involving antioxidants NADPH (***Fig. 1***), and GSH (***Fig. 2***), we carefully constructed a schematic of reactions and identified the known inhibitors of the redox components (***Fig. 3A***). Because our data suggest a positive correlation between enhanced anti-oxidation balance in melanoma cells and reduced drug sensitivity, we examined whether we could alter sensitivity of melanoma cells to BRAFi by using redox inhibitors. To evaluate the role of different redox components, we tested BRAF-inhibitors in combination with known redox inhibitors such as DPI, RSL3, Buthionine sulfoximine (BSO), Erastin, and FK866. We subjected two cell lines, one sensitive (WM88), and one insensitive (SC10) to increasing concentrations of either DPI, RSL3, or BSO alone or in combination with 8μM PLX4720 (***Fig. 3B***). While both cells exhibited concentration-dependent anti-proliferative effects to single redox inhibitors (DPI, RSL3, BSO), enhanced effects of combination were observed only in the insensitive, and not in cells already sensitive to PLX4720 (***Fig. 3B***). Similar effects were seen in A2058, considered as largely insensitive *BRAF*-mutated melanoma cells (***Fig. S2***). Depending on the drugs, the combination either increased efficacy (maximum effect), or potency (more effect with less drug) or both in drug-insensitive SC10. However, not all redox inhibitors enhanced the effects of PLX4720. Erastin (SLC7A11 inhibitor) alone did not exhibit any effects on SC10, and its combination with PLX4720 was similar to the effects of PLX4720 alone (***Fig. 3C***, black dotted line). Similarly, the combination of FK866 (which inhibits the enzyme NAMPT, and thus depletes NAD+) with PLX4720 was not advantageous compared to single agents alone. At lower concentrations, the effect of combination was similar to PLX4720 alone, and at higher concentrations, similar to FK866 alone, indicating that the combination did not enhance the effects of single agents (***Fig. 3C***). This is an interesting result, as it suggests that inhibition of only certain redox components may enhance the effects of BRAF-inhibitors in *BRAF*-mutated melanoma cells.

**Figure 3:**
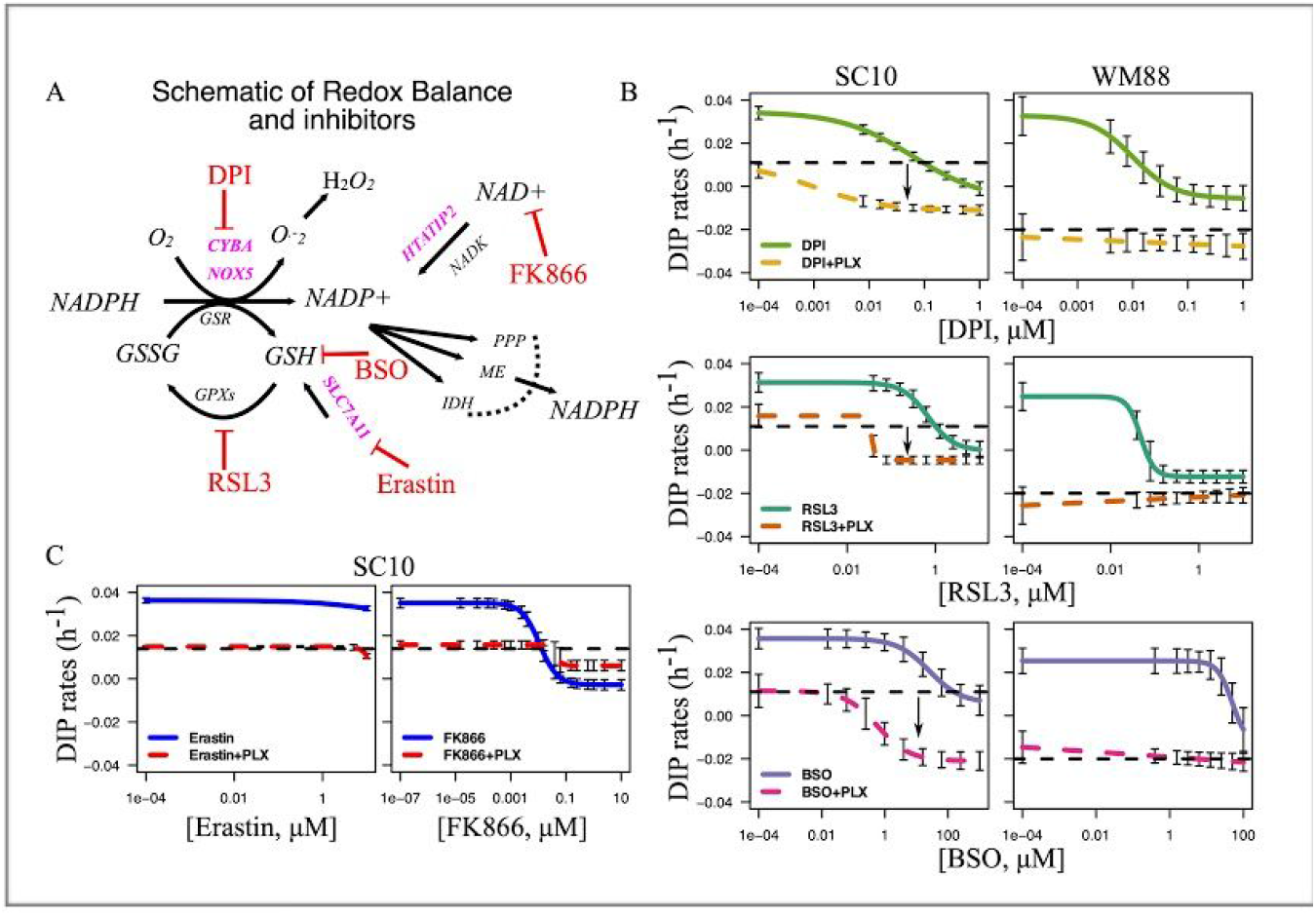
Pharmacological inhibition of redox enzymes enhances the effects of BRAFi in BRAF-mutated melanoma cells. (A) Schematic of reactions identified in Figure 1 with pharmacological inhibitors targeting each node as shown. (B) Drug-dose response curves of SC10 (left), and WM88 (right) either in single agents or in combination with 8μM PLX4720. Arrows show the enhanced effects of combinations, indicating that combination enhances the effects of single agents in drug-insensitive (SC10), and not in WM88 cells. (C) Drug-dose response curves of SC10 either in single agents or in combination with 8μM PLX4720. In both (B), and (C), black dotted lines represent the DIP rates in 8μM PLX4720; colored solid lines denote the DIP rates in increasing concentrations of single agents (DPI: NADPH Oxidase (NOX) inhibitor, RSL3: GPX4 inhibitor, BSO: GCLC inhibitor, potent depletion of GSH, Erastin: SLC7A11 inhibitor, FK866: NAD+ depletion, NAMPT inhibitor); while the colored dotted lines denote the drug responses of combination of 8μM PLX4720 and specific single agents corresponding to the colored solid curve (3+ biological replicates).

### Enhanced effects of combination are due to increased oxidative stress

Our drug combinations suggest that co-administration of certain redox inhibitors with PLX4720 enhance the benefits of targeted therapies in melanoma (***Fig. 3***). To investigate how co-administrations affect redox balance in melanoma cells, we stained *BRAF*-mutated melanoma cell, SKMEL5 treated with either single agents or combinations with CellROX® Deep Red Reagent. Specifically, we tested three drug combinations that showed increased drug effects with BRAF-inhibitors: DPI, RSL3, and BSO. Unlike in BSO, and RSL3, the total cellular reactive oxygen species (ROS) levels were increased upon PLX4720 treatment (p < 0.001) (***Fig. 4A*** ***& Fig. S3***), consistent with the results in earlier reports (Bauer et al., 2017). Although counter-intuitive, DPI treatment as single agent induced a significant increase in ROS intensity compared to DMSO control (***Fig. 4A*** ***& Fig. S3***). This result is consistent with studies that suggest DPI is a non-specific NOX-inhibitor, and causes an enhanced oxidative stress in cells (Riganti et al., 2004). DPI is a general flavoprotein inhibitor, and non-specific NOX-inhibitors; it is also shown to inhibit xanthine oxidase, eNOS etc (Altenhöfer et al., 2015). Similarly, all three drug combinations (BSO+PLX, RSL3+PLX, & DPI+PLX) resulted in a significant increase in total cellular ROS compared to DMSO control, PLX4720 alone or the respective single agents (***Fig. 4A*** ***& Fig. S3***). Our results reveal that co-administration of redox inhibitors with PLX4720 disrupts redox balance in cells, and causes an increase in oxidative stress. Therefore, we sought to test whether the lethal effects of these drug combinations could be mitigated by exogenous antioxidants. We excluded the combination of DPI and PLX4720, because of the broad off-target effects of DPI. For the other two combinations, we co-treated melanoma cells with two known antioxidants: reduced Glutathione (GSH), and lipophilic antioxidant Ferrostatin-1 (FER). The effects of PLX4720 alone could not be rescued by either GSH, and FER (***Fig. S4***), indicating that the antiproliferative effects of PLX4720 is not due to oxidative stress alone. Interestingly, supplementation of the culture medium with GSH substantially rescued both RSL3, and BSO-induced cell death in combination (***Fig. 4C***), while Ferrostain-1 rescue was specific to RSL3 only, confirming the glutathione levels as a mechanistic link for reduced drug sensitivity in *BRAF*-mutated melanoma cells.

**Figure 4:**
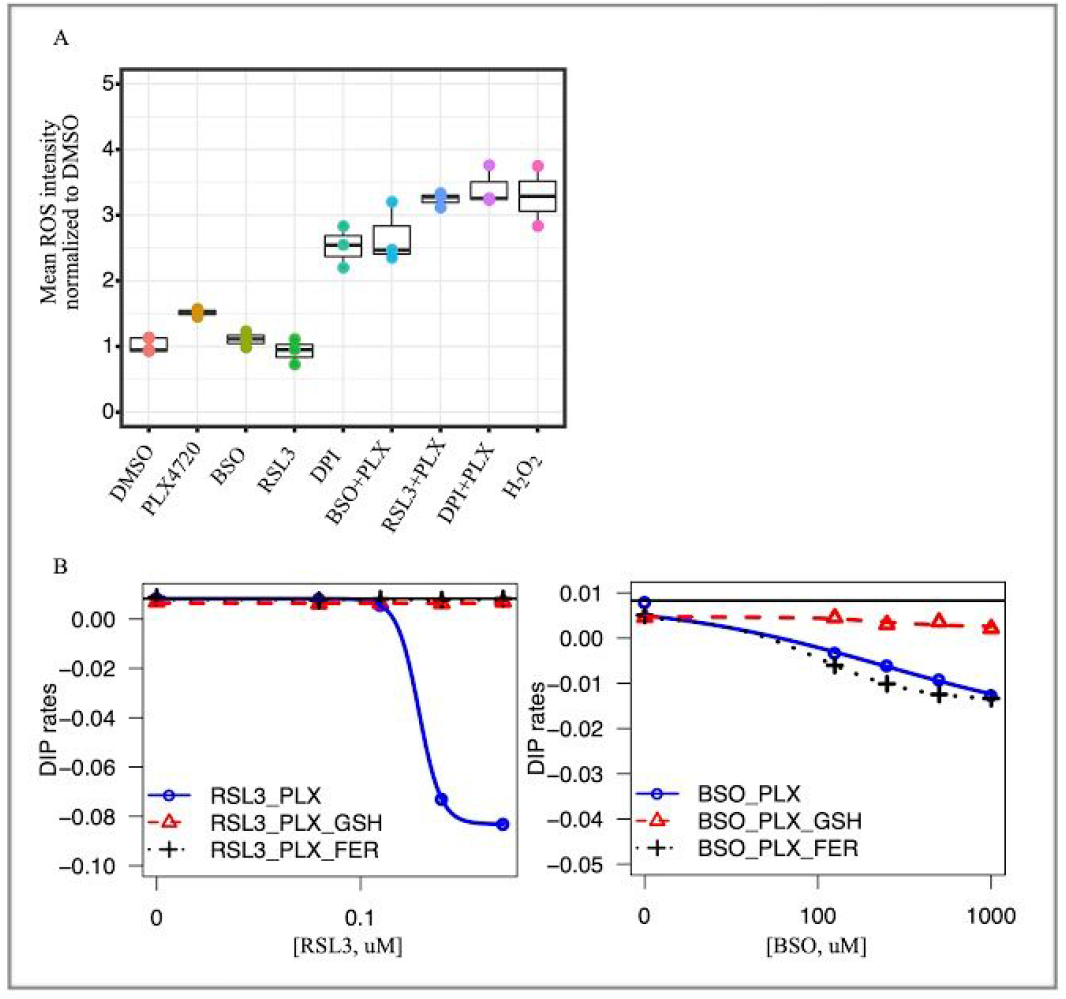
Co-treatment of redox inhibitors and BRAFi induces oxidative stress. (A) Box plots showing total cellular ROS mean intensity per cell for either single agents or in combinations normalized to intensity in DMSO control for melanoma cells (SKMEL5 fluorescently tagged with GFP-H2B) stained with CeIIROX® Deep Red Reagent. Cells were treated with DMSO control, 8μM PLX4720, and H_2_0_2_ (1mM incubated for an hour); middle panel has BSO (500μM), RSL3 (78.125nM), and DPI (1.25μM); in box plot, the solid line is the median, the box spans the first, and third quartiles, the whiskers extend to 1.5× the interquartile range. (B) Fitted drug-dose response curves of quantified DIP rates of SKMEL5 melanoma cells in drug combinations (RSL3+PLX, left), and combinations (BSO+PLX, right). Two known antioxidants--reduced glutathione (GSH, 5mM), and ferrostatin-1 (FER, 5μM) were added to the combinations at day 0, and replenished at 72 hrs along with growth medium, and inhibitor. Solid black lines denote the DIP rates in 8μM PLX4720. The effects can be rescued by GSH in both, while Ferrostatin-1 rescues only in RSL3 combination (3+ biological replicates).

### Pan-melanoma metabolic models exhibit altered redox metabolism and enhanced antioxidative capacity

While melanomas with reduced sensitivity to BRAF-inhibition show enhanced antioxidative capacity potentiating redox-targeting therapies, it remains unclear whether other melanoma subtypes display similar or varying redox phenotypes. To investigate whether alterations in redox metabolism are common attributes across melanomas, FBA models of 103 melanoma tumors from The Cancer Genome Atlas (TCGA) and 860 normal skin tissues from the Genotype-Tissue Expression (GTEx) project were developed using their respective gene expression profiles (Carithers and Moore, 2015; Lewis et al., 2018; Weinstein et al., 2013). Melanoma tumor models displayed greater steady-state fluxes through major NADPH-oxidizing reactions including GSR and DHFR compared to normal tissue models (***Fig. 5A***). Interestingly, normal tissue models predicted significantly greater flux through DUOX1/2, an NADPH-oxidizing enzyme involved in cellular ROS generation; thus, increased activity of antioxidant-generating GSR and decreased activity of ROS-generating DUOX1/2 likely contribute to the enhanced antioxidative capacity in melanoma tumors compared to normal tissues (Donkó et al., 2005). These findings are similar to the comparison in predicted metabolic fluxes between drug-insensitive SC10 cells and drug-sensitive SC01 cells (***Fig. 1F***), suggesting a possible progression in redox capacity from normal skin tissues (low capacity) to drug-sensitive melanomas (medium capacity) to drug-insensitive melanomas (high capacity). Using our FBA models of TCGA tumors, a simulated metabolome-wide gene knockout screen was performed, where for each of the 3,268 genes in the Recon3D metabolic reconstruction, the effect of gene knockout on total NADPH oxidation in each melanoma model was determined (***Fig. 5B***) (Brunk et al., 2018). Many of the gene knockouts with greatest effects on NADPH oxidation across TCGA melanoma models included major genes involved in the reduction of NADP+ to NADPH (e.g. GLUD1, GLUD2, IDH2, MTHFD1), as well as genes utilizing reduced NADPH to increase cellular antioxidant stores or decrease levels of reactive oxygen species (e.g. GSR, CBR1) (Hayes and Dinkova-Kostova, 2014; Lewis et al., 2018). Next, we sought to test whether our identified redox-targeting therapies work in other subtypes of melanoma. We applied the drug combinations (BSO+PLX, and RSL3+PLX) in *NRAS*-mutated (SKMEL2), and *NF1*-mutated (MeWo) melanoma cells. The combinations increased efficacy of BRAF-inhibitors in both SKMEL2, and MeWo cells, consistent with results above (***Fig. 5C***). Collectively, these findings suggest that altered redox metabolism and enhanced antioxidant production are common features among melanomas, and that inhibition of redox enzymes may be a viable treatment strategy for melanomas in general.

**Figure 5:**
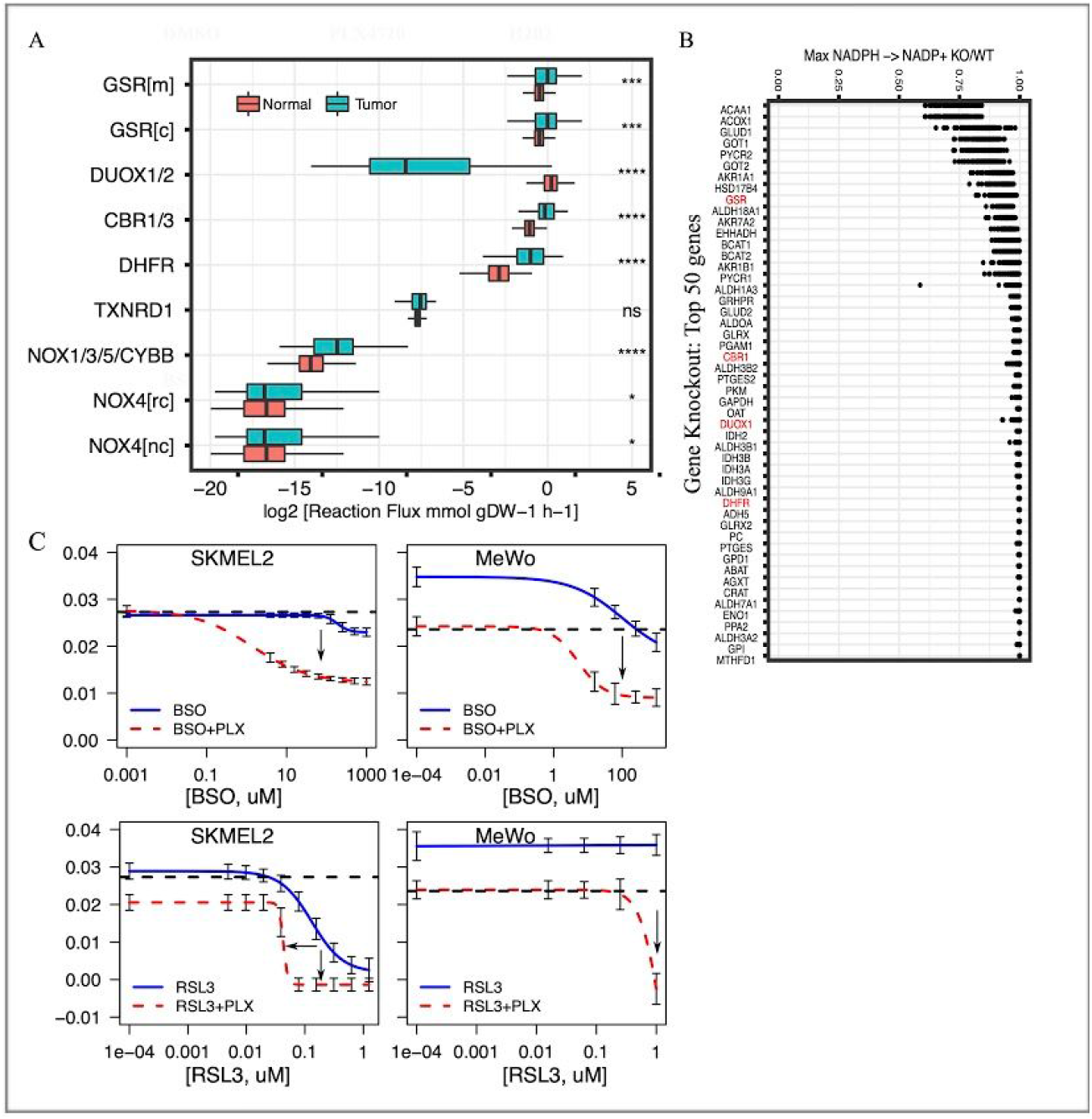
Average FBA predicted metabolic flux distributions in TCGA melanoma tumors compared to normal skin tissues identify key redox vulnerabilities in pan-melanoma tumors. (A) Box plots showing estimated average reaction fluxes through different enzymes in NADPH oxidation pathways in melanoma tumors (red box-plot) compared to normal skin (grey-green box-plot). Each dot represents a distinct tumor (expression from TCGA melanoma patient) or normal tissue (expression from Genotype-Tissue Expression (GTEx). *(B)* Effect of *in-silico* gene knockdown on predicted net conversion of NADPH to NADP+ in TCGA melanoma tumors, expressed as a ratio of net conversion after (KO) versus before knockdown (WT) (KO/WT). Shown are top 50 genes with greatest effects on NADPH net conversion, enzymes highlighted in red are genes in Fig. 5A. (C) Drug-dose response curves of NRAS-mutated melanoma cells (SKMEL2, left), and NF1-mutated melanoma cells (MeWo, right) in single agents or in combination with 8μM PLX4720. In both (left, and right), black dotted lines represent the DIP rates in 8pM PLX4720; colored solid lines denote the DIP rates in increasing concentrations of single agents (RSL3: GPX4 inhibitor, BSO: GCLC inhibitor, potent depletion of GSH); while the colored dotted lines denote the drug responses of combination of 8μM PLX4720 and specific single agents corresponding to the colored solid curve (3+ biological replicates). *p <= 0.05, ** p <= 0.01, *** p <= 0.001, **** p <= 0.0001

## Discussion

Overall, we report here that higher antioxidant ability of cells correlates to lower drug sensitivity to targeted therapies in melanoma. Our analyses reveal an association between an elevated redox capacity, involving two reducing cofactors NADPH and GSH, and reduced drug sensitivity in melanoma. Furthermore, we show that drug-insensitive melanoma cells not only possess a robust redox system, but also maintain key antioxidant metabolites during treatment. Indeed, we show that the disruption of this redox buffer improves the outcomes of targeted therapies. In summary, our findings suggest that modulation of cancer antioxidant defense could be exploited to augment the benefits of existing therapies in melanoma (Khamari et al., 2018; Yuan et al., 2018). This strategy may extend beyond melanoma, as antioxidant pathways have been implicated in tumor progression, and drug resistance (Harris et al., 2015; Ji et al., 2018; Sarmiento-Salinas et al., 2019).

How ROS affects cancer cells has been a contentious issue for years. ROS likely elicits a broad spectrum of cellular responses (Paudel and Quaranta, 2019). Although still debated, its effects on cancer cells can best be described as hormesis (Schieber and Chandel, 2014): at low levels, it can mediate sustained network signaling leading to enhanced proliferation (Sena and Chandel, 2012), while at high levels, it can induce cellular damage and cell death (Panieri and Santoro, 2016; Ray et al., 2012). The level of ROS that a cell can handle, thus, likely depends on how functional its antioxidant machinery is. In general, tumors can sustain a much higher levels of ROS compared to normal tissues by invoking their antioxidant systems, thereby creating a high redox homeostatic state (DeNicola et al., 2011). Melanoma tumors could possibly exist in such a high redox homeostatic state, as we see an increased activity of antioxidant-generating GSR, and decreased activity of ROS-generating DUOX1/2 in TCGA melanoma tumors compared to normal skin. However, there exists a significant variability even within tumors, captured well in melanoma cell lines, suggesting redox capacity perhaps could be on a continuum: low capacity (normal skin), moderate capacity (drug-sensitive melanomas), and high capacity (drug-insensitive melanomas).

Based on modeling by Flux Balance Analysis (FBA), the high redox capacity in drug-insensitive melanomas is predicted to stem from elevated fluxes through metabolic reactions which generate important antioxidant molecules including glutathione. This computational approach towards predicting genome-scale metabolic phenotypes has many advantages. By integrating the most comprehensive model of human metabolism to date with transcriptomic data from cell lines or tumors of interest, quantitative predictions on the metabolic status of these samples can be obtained, which would not be available from gene expression analysis alone (Blazier and Papin, 2012; Brunk et al., 2018; Lewis et al., 2018). Additionally, although FBA models can only make steady-state flux predictions, the interconnections between 13,000+ metabolic reactions in the human metabolic network often yield predictions with greatly improved accuracy and utility compared to smaller dynamic metabolic models which are missing these genome-scale interactions (Orth et al., 2010). Because of these advantages, FBA has led to many successful predictions in cancer metabolism, providing powerful insights into both cancer pathophysiology and treatment strategies (Asgari et al., 2015; Folger et al., 2011; Gatto et al., 2015; Lewis and Abdel-Haleem, 2013). Additionally, we used FBA to predict metabolic fluxes for reactions in central carbon metabolism (glycolysis and TCA cycle) in TCGA melanoma tumors, and compared them with GTEx normal skin tissues (***Fig. S5A***). Our analysis suggests that many reactions had a statistically significant difference between melanoma tumors and normal skin, with most glycolytic and some TCA cycle fluxes significantly increased in melanoma tumors (***Fig. S5A***). Interestingly, similar analyses between *BRAF*-mutated TCGA melanoma tumors compared with TCGA non *BRAF*-mutated melanomas show no difference in predicted metabolic fluxes through central carbon metabolism (***Fig. S5B***). By coupling FBA, gene expression, and metabolomics screen, we show that the drug-insensitive melanomas have an increased dependence on antioxidant system. This increased dependence could present a therapeutic opportunity in melanomas. Indeed, our own results and recent reports inform that redox modulation could be a viable strategy to improve the treatment outcomes (Bauer et al., 2017; Cesi et al., 2017; Corazao-Rozas et al., 2013; Fruehauf and Trapp, 2008; Gorrini et al., 2013; Yuan et al., 2018).

Glutathione (GSH) is the most abundant antioxidant in the cell. It can be synthesized either *de novo* or regenerated using NADPH as the substrate (Fan et al., 2014). *De novo* GSH biosynthesis requires cysteine, which is imported into the cells by cystine/glutamate antiporter, SLC7A11 (Ji et al., 2018; Shih and Murphy, 2001). Our data suggest that drug-insensitive melanoma cells have both upregulated SLC7A11 expression, and higher predicted fluxes through GSH regenerating GSR, suggesting an efficient maintenance of glutathione. Intriguingly, drug-insensitive melanoma cells can also maintain higher levels of GSH, and maximally utilize cysteine during treatment. In addition, the level of serine is higher in drug-insensitive cells, and increased upon treatment. Since serine is a precursor of cysteine, it may help cells maintain necessary levels of glutathione (Mattaini et al., 2016; Zhou et al., 2017). Therefore, we next questioned whether the modulation of GSH level affects drug sensitivity in melanoma. Inhibition of SLC7A11, however, did not improve the effects of BRAFi in melanoma cells, indicating that melanoma cells could maintain GSH levels through NADPH regeneration. In contrast, GSH depletion by BSO enhanced the effects of combination. Thus, GSH modulation appears as an exciting therapeutic target in melanoma. Efficient regeneration of GSH requires sufficient NADPH, another important cofactor in cells. Our work focussed on NADPH oxidation (consumption), and examined the contributions of several enzymes that convert NADPH to NADP+. Sixteen out of top 50 enzymes that had the most effect on the predicted fluxes through NADPH are known to be involved in NADPH production pathways. Examining the importance of NADPH production in melanoma drug sensitivity is a possible area of future investigation. We speculate that drug-insensitive melanoma cells may also possess a robust NADPH production pathways.

Consistent with earlier reports (Cesi et al., 2017; Ravindran Menon et al., 2015; Vazquez et al., 2013), we observed that BRAFi alone also induced ROS in melanoma. However, BRAFi induced effects on melanoma cells could not be rescued by known antioxidants, suggesting that effects of BRAFi on drug-insensitive melanoma cells are not solely due to ROS. Another possibility is that drug-insensitive melanoma cells perhaps utilize their robust antioxidant system to tolerate the induced ROS. Therefore, an increase in oxidative stress due to the combinations identified here, possibly disrupts that redox balance. One such agent for combination, RSL3, is known to induce ferroptosis in cancer cells. Our results are, thus, consistent with recent reports that suggest ferroptosis-inducing drugs may enhance the current treatment options for melanoma (Tsoi et al., 2018). Ferroptosis has recently been exploited in different cancer types as an alternative way to reduce the fraction of drug-tolerant persister cells (Hangauer et al., 2017; Viswanathan et al., 2017). Yet, one important consideration is that ROS is not one, but a family of chemical species. Therefore, it remains to be determined whether different species of ROS have different effects on melanoma cells. Further studies are required to ascertain the species of ROS that have detrimental effects on melanoma cells. One could imagine this could be cell-line or tumor specific, as gene regulatory network (GRN) has been linked with the ability of cells to undergo metabolic plasticity (Jia et al., 2019; Paudel and Quaranta, 2019). Future work seems necessary to also investigate the compensatory antioxidant pathways, and how they help maintain redox homeostasis in melanoma cells (Harris et al., 2015).

In summary, using an integrative approach, we identify new vulnerabilities in melanoma cells, that may extend beyond melanoma. We show that enhanced capacity of redox balance may provide an early survival advantage in melanomas against MAPK pathway inhibition. We propose that disrupting an antioxidant balance in melanomas offers a useful therapeutic target against cells with reduced sensitivity to existing therapies.

## Acknowledgments

We are grateful to Meenhard Herlyn (Wistar Institute), Kim Dahlman (Vanderbilt University), and Ann Richmond (Vanderbilt University) for kindly providing *BRAF*-, *NRAS*-, and *NF1*-mutated melanoma cell lines; to Joshua P. Fessel for insightful discussions on NADPH oxidases; to Darren R. Tyson, Christian T. Meyer, David J. Wooten, Leonard A. Harris for useful discussions; to Jing Hao for support in reagent procurement, and experimental preparation. We would also like to thank Prof. Kevin A. Janes (University of Virginia) for critical review of this manuscript.

This work was supported by the US National Institutes of Health Grants U54 CA217450, U01 CA215845, R01 CA186193, and U01 CA174706 (to V.Q.); U01 CA215848 (to M.L.K); Vanderbilt Institute for Clinical and Translational Research (VICTR) grants 16721, and 16721.1 (to B.B.P). J.E.L is supported by NIH/NCI F30 CA224968. C.E.H. is supported by NIH/NCI F31 CA221147. K.N.H. is now at Houston Methodist Research Institute, Houston, TX. C.J.R. is now at Yale University School of Medicine MD-PhD Program, New Haven, CT.

## Author Contributions

Conceptualization: B.B.P., and V.Q.;

Methodology: B.B.P., J.E.L., M.L.K., and V.Q.;

Investigation: B.B.P., J.E.L., K.N.H., C.E.H., C.J.R., G.S.C., S.D.S.;

Formal Analysis: B.B.P., J.E.L., G.S.C., S.D.S., J.A.M., M.L.K., and V.Q.;

Writing-Original Draft: B.B.P., J.E.L.;

Writing-Review & Editing: B.B.P., J.E.L., K.N.H., G.S.C., S.D.S., J.A.M., M.L.K., and V.Q.;

Visualization: B.B.P., J.E.L.;

Supervision: J.A.M., M.L.K., and V.Q.;

Funding Acquisition: M.L.K., and V.Q.

## EXPERIMENTAL MODEL AND SUBJECT DETAILS

All *BRAF*-mutated, *NRAS*-mutated, *NF1*-mutated melanoma cells were engineered to express either histone 2B-mRFP using pHIV-H2B-mRFP plasmid (Welm *et al.*, 2008) and/or Fluorescent Ubiquitination Cell Cycle Indicator (FUCCI) using Geminin1-110 monomeric Azami Green plasmid (Karasawa et al., 2003) as previously described (Tyson et al., 2012). Single-cell derived clonal populations of BRAF-mutated melanoma cells, SKMEL5 were selected by limiting dilution as previously described (Paudel et al., 2018).

## METHOD DETAILS

### Cell Culture and Chemical Reagents

Single-cell derived *BRAF*-mutated SKMEL5 subclones were derived as previously described (Paudel et al., 2018). *BRAF*-mutated melanoma cells (SKMEL5, WM88), including the SKMEL5 subclones, NRAS-mutated melanoma cells (SKMEL2), and NF1-mutated melanoma cells (MeWO) were grown and cultured in Dulbecco’s modified Eagle’s medium and Ham’s F-12 media (DMEM:F12, 1:1, Cat. No. 11330-032). Media were obtained from Gibco (Grand Island, NY), and supplemented with 10% fetal bovine serum. All cells were cultured in humidified incubators that were CO2 and temperature (37°C) controlled. Cells were passaged 1–2 times per week and were maintained as exponentially growing cultures for a maximum of less than 20 passages. All cells were tested for mycoplasma, and tested negative.

PLX4720 (Cat. No. S1152) and vemurafenib (Cat. No. S1267) were obtained from Selleckchem (Houston, TX). Dabrafenib (Cat No. HY-14660), (1S,3R)-RSL3 (Cat No. HY-100218A), Ferrostatin-1 (FER1, Cat No. HY-100579), Erastin (Cat No. HY-15763), (E)-Daporinad (FK866) (Cat No. HY-50876) were obtained from MedChem Express (Monmouth Junction, NJ) and solubilized in dimethyl sulfoxide (DMSO) at a stock concentration of 10 mM. Powdered L-Buthionine-sulfoximine (BSO) (Product No. B2515), powdered Diphenyleneiodonium chloride (DPI) (Product No. D2926), and powdered L-Glutathione reduced (GSH) were obtained from Sigma-Aldrich. BSO, and GSH was freshly made at a stock concentration of 100 mM in H_2_O, while DPI was solubilized in DMSO at a stock concentration of 10mM. CellRox™ DeepRed Reagent (Cat No. C10422) for oxidative stress detection was obtained from ThermoFisher Scientific. Acetonitrile (AcCN) (Cat No. A955-1), Methanol (MeOH) (Cat No. A456-1) and water (Cat No. W6-1), Optima LC-MS grade, for the mass spectrometry analysis were obtained from ThermoFisher Scientific.

### Cellular ROS Staining Assay and Quantification

CellRox™ DeepRed Reagent was used according to the manufacturer’s instructions. Briefly, melanoma cells were seeded in 96-well plates and treated with inhibitors for a duration specified. Hydrogen Peroxide (H_2_O_2_) was used as a positive control. Cells were incubated with 1mM H_2_O_2_ mixed with growth media for an hour before incubation with CellRox reagent. The cells were then stained with 5uM CellRoxTM DeepRed reagent by diluting the probe in complete growth media and incubated for 30 minutes at 37°C in tissue-culture incubators. Cells were then washed with PBS three times and imaged through a 20x objective with a Cellavista HighEnd Bioimager (SynenTec Bio-Services, Munster, Germany). Total ROS intensity was quantified by image segmentation in Fiji, image processing package (Schindelin et al., 2012).

### Bioinformatics Analysis Pipeline

RNASeq of melanoma cell lines (SKMEL5 subclones) was performed as previously reported (Meyer et al., 2019). Transcript count data from Gene Expression Omnibus (Accession number: GSE122041, https://www.ncbi.nlm.nih.gov/geo/query/acc.cgi?acc=GSE122041) was used for downstream analysis performed in R (https://www.r-project.org). Differentially Expressed Genes (DEGs) were selected by ANOVA on baseline gene expression on three SKMEL5-derived subclones based on a statistical cutoff of Likelihood Ratio Test (LRT) (false discovery rate (FDR) < 0.001, log2-fold change (LFC) >1). Differential expression analyses were performed using Bioconductor (https://www.bioconductor.org) packages DESeq2 (Love et al., 2014) and edgeR (Robinson et al., 2010). Genes present in results from both pipelines: DESeq2, and edgeR (common genes) were selected for further analyses. Of total 2165 DEGs, 1361 were upregulated and 804 were downregulated on a pairwise comparison between SC10 and SC01. Expressions of the identified DEGs were correlated against BRAFi sensitivity measured as DIP rate at 8uM PLX4720 (Hardeman et al., 2017) for 10 melanoma cell lines in CCLE database (Barretina et al., 2012). Only genes with Pearson’s correlation score of 0.5, and p < 0.05 were selected as significantly correlated genes (SCGs). Of total 71 SCGs, 36 were positively correlated, while 35 were negatively correlated. Functional Gene enrichment analyses were performed using clusterProfiler (Yu et al., 2012), and returned Gene Ontology (GO) terms were summarized using REVIGO (Supek et al., 2011).

### Flux Balance Analysis (FBA)

FBA is a computational approach for predicting genome-scale steady-state metabolic fluxes in samples of interest (Orth et al., 2010). Reaction fluxes throughout the metabolic network are predicted by solving the optimization problem:

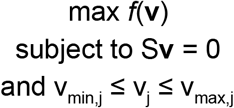

where if *m* and *r* are the number of metabolites and reactions in the metabolic network, respectively, then **v** is a *r* × 1 vector of reaction fluxes to be solved for, S is the *m* × *r* stoichiometric matrix of the metabolic network, v_min,j_ and v_max,j_ are constraints imposed on each individual reaction flux, and *f* is an objective function that is maximized to maximize a particular metabolic phenotype of interest. Recon3D version 3.01 was used as the core metabolic network (Brunk et al., 2018).

To maximize the oxidation of NADPH to NADP^+^ in the metabolic network, the following objective function was used in all cellular compartments:

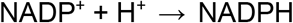

This artificial sink on NADP^+^ in turn maximizes the flux through existing metabolic reactions that oxidize NADPH to NADP^+^. Individual reaction flux values were predicted by averaging the minimum and maximum predicted fluxes from Flux Variance Analysis (FVA), which computes the minimum and maximum obtainable flux for each reaction under the model constraints and while maximizing the objective function above.

Personalized metabolic models for cell lines, tumors, and normal tissues were developed using transcriptomic data from these respective samples as previously reported (Lewis et al., 2018).

### Drug Combination Assay

The dose-response curves are generated using a 2-fold dilution of single drugs at concentrated indicated (highest) to zero (solvent the drug was dissolved in, either DMSO or H_2_O), all concentrations contained equal percentage of the solvent used. For combination with PLX4720, 8μM PLX4720 was used as diluant, and serially diluted for second drug from highest concentration to lowest. Direct measurements of cell counts were made using Cellavista software and ImageJ macros as previously described (Hardeman et al., 2017; Harris et al., 2016; Paudel et al., 2018). The drug-induced proliferation (DIP) rates are calculated using the slope of the log2-normalized population curves after 24 hrs.

### Antioxidant Rescue Assay

For drug-combinations indicated, the dose-response curves are generated as described above. For the indicated concentrations, two known antioxidants--reduced glutathione (GSH, 5mM), and ferrostatin-1 (FER, 5μM) are added to the combinations at day 0, and replenished at 72 hrs along with growth medium, inhibitors combination for the duration of the assay upto 6 days.

### Metabolomics Analysis

#### Sample Preparation

The combined global untargeted-targeted metabolomic analysis used BRAF-mutated melanoma cells (WM88, SKMEL5-SC10) treated with either DMSO or 8μM PLX4720 for 24 hrs. Cell pellet samples were lysed using 400μL ice cold lysis buffer (1:1:2, ACN:MeOH:Ammonium Bicarbonate 0.1M, pH 8.0, LC-MS grade) and vortexed well until the cells mixed well with the solvent. Each sample was sonicated using a probe tip sonicator, 10 pulses at 30% power, cooling down in ice between samples. A BCA protein assay was used to determine the protein concentration for each individual sample, and adjusted to a total amount of protein of 200μg total protein in 200 μL of lysis buffer. Heavy labeled standard molecules, Phenylalanine-D8 (CDN Isotopes, Quebec, CA), and Biotin-D2(Cambridge Isotope Laboratories, Inc., MA, USA), were added to each sample to assess sample handling steps. Samples were subjected to protein precipitation by addition of 800μL of ice cold methanol (4x by volume), then incubated at −80°C overnight. Samples were centrifuged at 10,000 rpm for 10 min to eliminate precipitated proteins and the metabolite containing supernatant was dried *in vacuo* and stored at −80°C until LC-MS analysis.

#### Global untargeted-targeted LC-MS/MS analysis

For mass spectrometry analysis, metabolite extracts were reconstituted in 60μl of HILIC resuspension buffer (AcCN/H_2_O, 90:10, v/v) and cleared by centrifugation. Quality control (QC) samples were prepared by pooling equal volumes from each sample. Stable isotope labeled standards, Tryptophan-D3, Carnitine-D9, Valine-D8, and Inosine-4N15, were added to each sample to assess MS instrument reproducibility.

Parallel reaction monitoring (PRM) was performed on a high resolution Q-Exactive HF hybrid quadrupole-Orbitrap mass spectrometer (Thermo Fisher Scientific, Bremen, Germany) equipped with a Vanquish UHPLC binary system and autosampler (Thermo Fisher Scientific, Bremen, Germany). PRM was used to monitor 18 metabolites with the following parameters: MS scan at 60,000 resolution with an automatic gain control (AGC) of 1e6, max injection time (IT) of 100 ms, and scan range from *m/z* 70–1050. Following the MS scan, six MS/MS scans were acquired at a resolution of 15,000, AGC value of 1e5, max IT of 50 ms,1.3 *m/z* isolation window and a stepped normalized collision energy. The targeted-MS^2^ (or PRM) method is a scheduled (± 2 min)inclusion list for the mass-to-charge ratios(*m/z)* of the metabolites of interest. These data are empirically derived based on previous LC-MS/MS analysis of pure (> 90%) standards.

MS and MS/MS data-dependent acquisition (DDA) were also performed on the Q-Exactive HF using the QC sample. Metabolite extracts (5uL injection volume) were separated on a SeQuant ZIC-HILIC 3.5-μm, 2.1 mm × 100 mm column (Millipore Corporation, Darmstadt, Germany) held at 40°C. Liquid chromatography was performed at 200 μl min^−1^ using solvent A (5mM Ammonium formate in 90% water, 10% acetonitrile) and solvent B (5mM Ammonium formate in 90% acetonitrile, 10% water) with the following gradient: 95% B for 2 min, 95-40% B over 16 min, 40% B held 2 min, and 40-95% B over 15 min, 95% B held 10 min (gradient length 45 min).

MS analyses were acquired over a mass range of m/z 70-1050 using electrospray ionization positive mode. MS scans were analyzed at a resolution of 120000 with a scan rate of 3.5 Hz. The AGC target was set to 1 × 10^6^ ions, and maximum ion IT was at 100 ms. Source ionization parameters were optimized, these include: spray voltage - 3.0 kV, transfer temperature – 280 °C; S-lens - 40; heater temperature - 325 °C; sheath gas - 40, aux gas - 10, and sweep gas flow - 1.

Tandem spectra were acquired using a data dependent acquisition in which one MS scan is followed by 2, 4 or 6 MS/MS scans. MS/MS scans are acquired using an isolation width of 1.3 *m/z*, stepped NCE of 20 and 40, and a dynamic exclusion for 6 s. MS/MS spectra were collected at a resolution of 15000, with an AGC target set at 2 × 10^5^ ions, and maximum ion IT of 100 ms. Instrument performance and reproducibility in the run sequence was assessed by monitoring the retention times and peak areas for the heavy labeled standards added to the individual samples prior to and during metabolite extraction to assess sample processing steps and instrument variability(***Table S7***).

#### Metabolite data processing

LC-MS/MS raw data were imported, processed, normalized and reviewed using Progenesis QI v.2.1 (Non-linear Dynamics, Newcastle, UK). All MS, DDA and PRM sample runs were chromatographically aligned against a QC reference run. Following peak picking, unique spectral features (retention time and m/z pairs) were grouped based on adducts and isotopes, and individual features or metabolites were normalized to all features. Further filtering was carried out by removing features or metabolites that had >30% coefficient of variance.

## QUANTIFICATION AND STATISTICAL ANALYSIS

Data is presented as the *mean* ± *SD* or *mean* ± *SEM* where appropriate from at least 3 biological replicates, unless stated otherwise in figure legends. Statistical analysis was performed as indicated in figures, figure legends, or experimental methods in R. Data analysis was performed in the R statistical analysis software package, R version 3.5.1, and R-studio version 1.1.463 (https://www.r-project.org/ and https://www.rstudio.com/).

## DATA AND CODE AVAILABILITY

All data, calculated DIP rates, DEGs between subclones, SCGs for CCLE melanoma panel, expression data, codes used to generate the figures are available in the GitHub repo: GitHub Repo.

## Supplemental Information

Supplemental information can be found at: Supplementary Info.

